# Sleep-like Slow Waves During Wakefulness Mediate Attention and Vigilance Difficulties in Adult Attention-Deficit/Hyperactivity Disorder

**DOI:** 10.1101/2025.07.27.666103

**Authors:** Elaine Pinggal, James Jackson, Anikó Kusztor, David Chapman, Jennifer Windt, Sean P.A. Drummond, Tim J. Silk, Mark A. Bellgrove, Thomas Andrillon

**Affiliations:** School of Psychological Sciences, Monash University, Clayton, VIC, 3800, Australia; Turner Institute for Brain and Mental Health & School of Psychological Sciences, Faculty of Medicine, Nursing, and Health Sciences, Monash University, Melbourne, VIC, 3800, Australia; Monash Alfred Psychiatry Research Centre (MAPrc), Melbourne, VIC, 3004, Australia; School of Philosophical, Historical, and International Studies, Monash University, Melbourne, VIC, 3800, Australia; Monash Centre for Consciousness and Contemplative Studies, Monash University, Melbourne, VIC, 3800, Australia; Centre for Social and Early Emotional Development (SEED) and School of Psychology, Deakin University, Burwood, Australia; Developmental Imaging, Murdoch Children’s Research Institute, Melbourne, Australia; Paris Brain Institute, Sorbonne Université, Institut National de la Santé et de la Recherche Médicale-Centre National de la Recherche Scientifique, Paris 75013, France

**Author notes:** These authors contributed equally. **Corresponding author:** Elaine Pinggal,; Thomas Andrillon. **Declaration of Conflict of Interests/Funding**. Authors have no Conflict of Interest to declare. This research was supported by a NHMRC Ideas Grant (“LAPSE Study”, APP2002454) and an ERC Starting Grant (“SleepingAwake”, 101116748).

**Keywords:** Electroencephalogram, attention-deficit/hyperactivity disorder, slow waves, attention, mental states, arousal

## Abstract

**Background:** Attention-Deficit/Hyperactivity Disorder (ADHD) is characterised by behavioural variability and heightened inattention associated with increased mind wandering (MW) and mind blanking (MB). Individuals with ADHD frequently experience sleep disorders and excessive daytime sleepiness, suggesting interactions between attention and arousal systems. Research examining brain activity using electroencephalography (EEG) has demonstrated that sleep-like slow waves (SW) during wakefulness are linked to inattention in neurotypical individuals, particularly following sleep deprivation, yet their role in ADHD remains unclear. This study investigated whether individuals with ADHD present with altered waking SW distribution compared to neurotypical controls and whether SW explain attentional difficulties in ADHD.

**Methods:** Adults with (*n* = 32) and without ADHD (*n* = 31) completed a sustained attention task while EEG recorded brain activity. Mental state probes (on-task, MW, MB) were embedded within the task. Sleep-like SW reflect a slowing of cortical activity and was detected from EEG activity. Omission/commission errors, reaction time (RT), RT variability, mental state reports and subjective sleepiness were analysed. Mediation analysis examined whether SW density explained ADHD-related performance differences.

**Results:** Individuals with ADHD exhibited more commission errors, MW and MB, and higher SW density (SW/min), particularly over parieto-temporal electrodes. Increased SW density correlated with higher omission errors, slower RTs, greater RT variability, and elevated sleepiness ratings. On-task reports were negatively correlated with SW density. Mediation analysis revealed that SW density significantly accounted for ADHD-related attentional difficulties.

**Conclusions:** Wake SW may explain attentional difficulties in ADHD, providing a potential mechanistic link between sleep disturbances and attentional fluctuations.

## Introduction

Attention Deficit/Hyperactivity Disorder (ADHD) is a common neurodevelopmental condition affecting approximately 2.5% of adults (Faraone et al., 2024). ADHD is characterised by excessive levels of inattention, hyperactivity and/or impulsivity (Faraone et al., 2024; May et al., 2023) which impact cognitive performance, particularly in tasks requiring sustained attention. Studies utilising sustained attention tasks have consistently demonstrated that individuals with ADHD exhibit greater response time variability and higher rates of commission and omission errors compared to neurotypical individuals, indicative of a fundamental difficulty with top-down attentional control (Bellgrove et al., 2005; Bozhilova et al., 2020, 2021; Tucha et al., 2017). Alongside these cognitive difficulties, individuals with ADHD also report an increased frequency of mind wandering (MW; operationally defined here as thinking about something else than the task at hand) as measured through both self-report questionnaires (Mowlem et al., 2019; Seli et al., 2015) and experience sampling methods during cognitive tasks (Bozhilova et al., 2021; Madiouni et al., 2020). These results emphasise the importance of considering subjective experience in ADHD in addition to cognitive performance.

MW is a common occurrence in the general population, with some estimates suggesting it occurs in up to 50% of waking hours (Killingsworth & Gilbert, 2010). MW is typically assessed during cognitive tasks by contrasting it with task-focused states based on the implicit assumption that individuals are continuously engaged with some form of conscious thought – either task-related or unrelated. However, recent findings show that people can also reliably report episodes of mind blanking (MB) (Andrillon et al., 2025), which are subjectively characterised by a momentary absence of conscious thought (i.e., thinking about nothing). When given the option to report both MW and MB, children and adults with ADHD reported more frequent MB, at the expense of both MW and task-focused states (Van Den Driessche et al., 2017).

Both MW and MB are associated with decreased behavioural performance (e.g., more omission and/or commission errors depending on the task), and represent the subjective side of attentional lapses (Smallwood & Schooler, 2015). Yet MW and MB can be associated with distinct behavioural signatures: in a Sustained Attention to Response Task (SART) (Robertson et al., 1997), MW is typically associated with impulsive responses, whereas MB is associated with sluggish responses (Andrillon et al., 2021). Additionally, both states have been associated with lower arousal as measured with pupil size or subjective ratings, suggesting a relationship between arousal and inattention over time. This has particular relevance in ADHD research, where dysregulated arousal and attentional variability are core features (Faraone et al., 2024; May et al., 2023).

Individuals with ADHD report disproportionately higher rates of sleep disturbances, including insomnia, parasomnias and increased daytime sleepiness (Bioulac et al., 2015; Bjorvatn et al., 2017; Hvolby, 2015). This pattern of disrupted sleep and excessive daytime sleepiness is associated with elevated electroencephalography (EEG) frontal delta activity during rest (Helfer et al., 2020). Since delta activity is typically observed in sleep using EEG, particularly over frontal cortices, its presence during wakeful rest in individuals with ADHD may reflect sleep-like activity within wakeful states, a phenomenon known as “local sleep” (Andrillon et al., 2019; Vyazovskiy et al., 2011). These sleep-like activity in wake manifest as high-amplitude, sleep-like slow waves, and have been associated with brief episodes of decreased neuronal activity in human and animal studies (Steriade, 2005; Vyazovskiy et al., 2011). These sleep-like slow waves increase with time spent awake or time spent on task, and have been linked to fluctuations in sustained attention in the general population (Andrillon et al., 2021; Bernardi et al., 2015; Hung et al., 2013). Importantly, the behavioural effects of these slow waves vary according to where they occur on the scalp, with frontal slow waves being associated with impulsive behaviour and increases in MW and MB (Andrillon et al., 2021; Pinggal et al., 2022), potentially reflecting the frontal region’s role in executive functions. In contrast, slow waves over parietal regions are associated with cognitive lapses (slower responses or misses), aligning with the parietal cortex’s involvement in sensory integration and decision-making (Freedman & Ibos, 2018). Thus, sleep-like slow waves during wakefulness could account for the varied cognitive profiles of individuals with ADHD (increased MW, MB, impulsive and sluggish responses, and excessive daytime sleepiness) and result from the sleep disturbances associated with ADHD.

Building on these neurophysiological findings, the present study sought to investigate whether slow wave activity during wakefulness was more prevalent in individuals with ADHD and whether this occurrence accounted for their attentional fluctuations compared to neurotypical individuals. To examine attentional difficulties at the behavioural and experiential level, we used a Sustained Attention to Response Task (SART) with integrated thought probes, prompting participants to report their current mental state. These recordings were paired with high-density EEG to capture slow wave activity. We aimed to examine behavioural and neural differences between the groups. Our primary hypotheses were that the ADHD group would: (1) demonstrate decreased task-performance, (2) report more frequent MW and MB (3) exhibit increased sleep-like slow waves relative to the neurotypical group. We also hypothesised that this increase in slow wave activity would mediate the effect of ADHD on task performance and subjective experience.

## Methods and Materials

### Participants

Sixty-five (*N* = 65; 33 with ADHD and 32 neurotypical individuals) participants were recruited through online advertisements, word of mouth, and flyers around Monash University. Neurotypical participants were screened to ensure they had no history of neuropsychiatric conditions, and their *t*-scores on subscales G and H of the Conners’ Adult ADHD Rating Scales – Self Report: Long Version (CAARS–S:L) were below 60 (indicating that they did not meet criteria for ADHD). Two participants were excluded from analyses: one participant due to data corruption and another due to misunderstanding task instructions. The final sample consisted of 63 participants (N = 32 ADHD and 31 neurotypical adults; see Table 1 for demographics and clinical characteristics). Except in one case, clinical interviews (Diagnostic Interview for ADHD in Adults (DIVA 2.0)) were conducted by a trained clinician to confirm ADHD diagnoses and determine the patients’ ADHD presentation: “inattentive” (N = 9), “hyperactive/impulsive” (N = 0) or “combined” (N = 22). Participants completed online questionnaires assessing demographics, ADHD symptoms (Conners’ Adult ADHD Rating Scales – Self Report: Long Version (CAARS), Adult ADHD Self-Report Scale for DSM-5 (ASRS-5), Wender Utah Rating Scale), sleepiness (Epworth Sleepiness Scale (ESS)), depression and anxiety (subscales from the Inventory of Depression and Anxiety Symptoms (IDAS-II)), mind wandering (part 1 of the Imaginal Process Inventory), chronotype (Morningness-Eveningness Questionnaire), and substance use (Alcohol Use Disorders Identification Test, Drug Use Disorders Identification Test). The results from the DIVA, CAARS and ESS questionnaires are presented in Table 1, while findings from the remaining measures are provided in Supplementary Table A. Participants with ADHD ceased medication

**Table 1.** Demographics and Questionnaire Data.

|  | ADHD (*N* = 32) | Neurotypical (*N* = 31) | *p*-Value |
| --- | --- | --- | --- |
| Age (years, mean (SD)) | 23 (3.5) | 23 (3.8) | .736 |
| Sex (% females) | 93.8% | 77.4% | .064 |
| Education level (years, mean (SD)) | 15.8 (1.8) | 15.5 (2.4) | .592 |
| Other lifetime psychiatric diagnoses |  |  |  |
| Depression | 50% | 3.2% |  |
| Any Anxiety Disorder | 62.5% | 0% |  |
| Personality Disorder | 3.13% | 0% |  |
| Autism | 21.9% | 0% |  |
| Post Traumatic Stress Disorder | 3.13% | 0% |  |
| Eating Disorder | 3.13% | 0% |  |
| Obsessive Compulsive Disorder | 3.13% | 0% |  |
| Questionnaire data |  |  |  |
| **Conners' Adult ADHD Rating Scale (CAARS)** |  |  |  |
| DSM-IV Inattentive Symptoms (Subscale E, mean (SD)) | 80.3 (8.5) | 50.8 (9.8) | < .001 |
| DSM-IV Hyperactive-Impulsive Symptoms (Subscale F, mean (SD)) | 65.9 (11.3) | 43.0 (7.9) | < .001 |
| DSM-IV Total ADHD Symptoms (Subscale G, mean (SD)) | 77.0 (8.2) | 47.6 (9.7) | < .001 |
| ADHD Index (Subscale H, mean (SD)) | 72.1 (9.1) | 45.5 (9.4) | < .001 |
| **Epworth Sleepiness Scale (mean, (SD))** | 8.7 (4.6) | 6.9 (3.6) | .098 |
| ADHD presentation (DIVA diagnosis) |  |  |  |
| Combined | 70.97% |  |  |
| Inattentive | 29.03% |  |  |
| Hyperactive-impulsive | 0% |  |  |

**Note.** ADHD = Attention Deficit/Hyperactivity Disorder. Higher scores on the CAARS subscales indicate greater severity of ADHD-related symptoms. Higher scores on the ESS indicate greater level of daytime sleepiness.

72 hours before testing (details of participants’ ADHD medication are provided in Supplementary Table B). The protocol was approved by Monash University Human Research Ethics Committee (project number: 17839).

### Modified Sustained Attention to Response Task (SART)

Participants completed a modified digit SART (Fig. 1a), in which pseudo-random sequences of digits (1–9) were presented on a screen. Stimuli were presented via Psychtoolbox (version 3) in MATLAB (Mathworks, Natick, MA, USA) with an inter-stimulus interval of 1-1.25s (random uniform jitter). Participants were instructed to respond with a button press (Go trials) every time they saw a new digit except for the digit 3 (No-Go trials).

**Figure 1.**
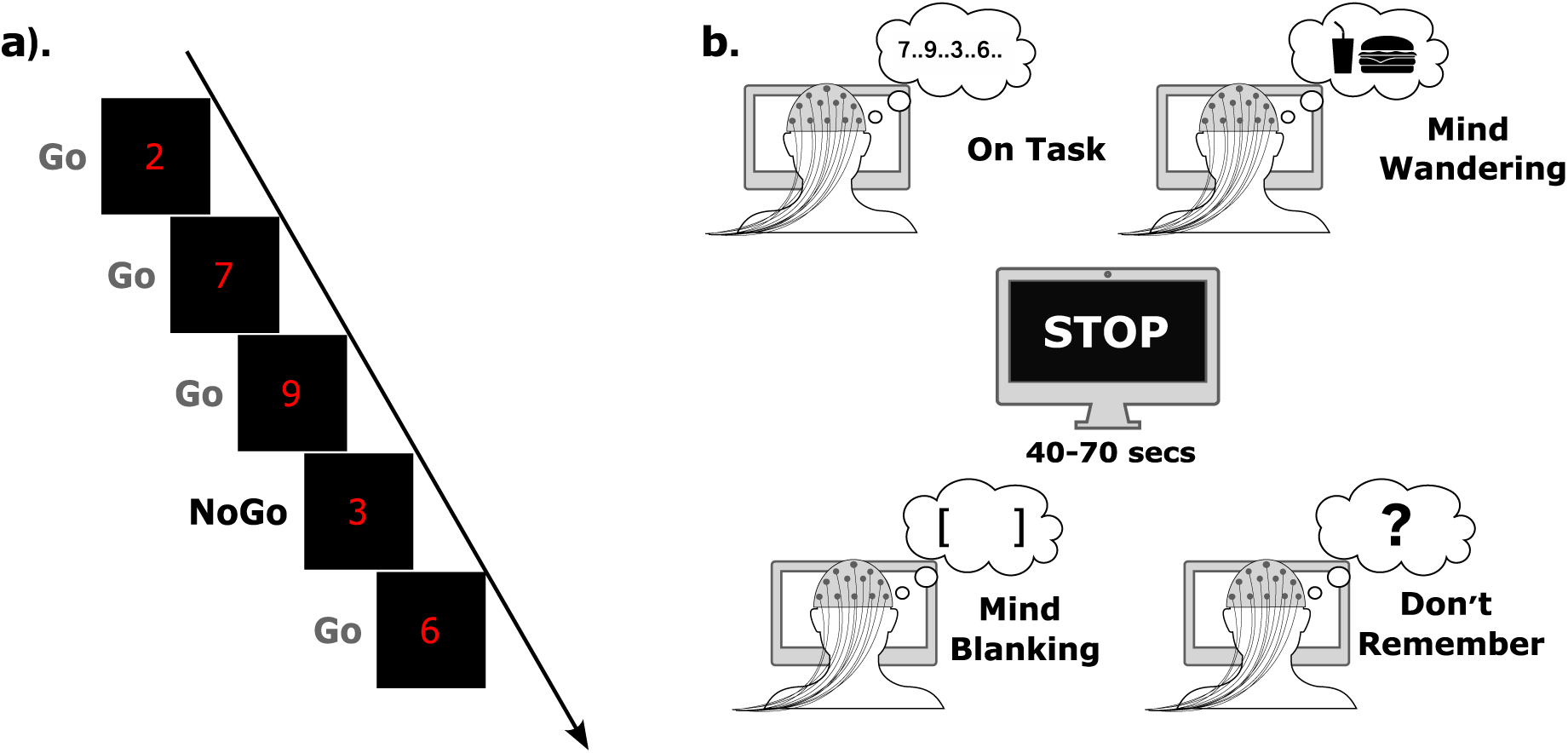
Experimental protocol. **a.** Participants completed four blocks of a modified SART where they were instructed to respond to every number except the number 3. **b.** Every 40–70 secs, participants were asked a series of 3–4 questions about their mental state immediately prior to the interruption.

Every 40–70s, participants were interrupted and asked a series of questions regarding their attentional focus (mental state) and alertness (Fig. 1b). Participants were first asked: *"Where was your attention focused? On-Task: Press 1, Off-Task: Press 2, Blank: Press 3, Don’t Remember: Press 4"*, where the response options corresponded to (1) thinking about the SART, (2) thinking about something other than the SART, (3) thinking about nothing, or (4) don’t remember. In cases where participants answered being “*off-task*” (option 2), they were subsequently asked: *“What distracted your attention from the task? Something:* (1) *in the room*, (2) *personal*, (3) *about the task*”. All participants were then prompted to rate the intentionality of their mental state with the question: *“Was your state of mind intentional? From 1: entirely intentional to 4: entirely unintentional”.* Finally, participants rated their alertness using the question: *“How alert have you been: Extremely alert: Press 1, Alert: Press 2, Sleepy: Press 3, Extremely sleepy: Press 4”*.

The task lasted for 52 (± 0.3 SEM across participants) minutes and consisted of four blocks of 12-15 minutes with 10 probes for each block (total: 40 probes and an average of 2120 trials). The main task was preceded by a 27-trial training session, with three digit (1–9) pseudorandomised sequences.

### Behavioural Data Analysis

For each trial, we defined (1) omission errors as the absence of a response during a Go trial, (2) commission errors as a response during a No-Go trial (digit 3), (3) reaction times (RTs) as the time between the digit presentation and the participant’s response on correctly detected Go trials, and (4) RT variability as the intra-individual coefficient of variation (ICVRT). Omission errors reflect a failure to respond to relevant visual stimuli, whereas commission errors indicate a failure to inhibit a response. Trials with RTs shorter than 150 ms were excluded from analyses. In addition, we computed the *d*-prime, a sensitivity index reflecting the ability to discriminate Go from No-Go trials, and criterion, a response bias measure indicating the tendency to favour Go or No-Go responses.

From the integrated thought probes, we extracted the percentage of each reported attentional state (on task, MW, MB, *“don’t remember”*), as well as average intentionality ratings for MW and average sleepiness ratings for each group.

We extracted behavioural performance metrics, attentional states and sleepiness ratings from a 20s window preceding each probe, as in Andrillon et al. (2021), ensuring the inclusion of two No-Go trials while capturing trials temporally close to probe presentation (Supp. Fig. 1). The behavioural patterns observed in these pre-probe windows were consistent with the results from our main behavioural analysis that included all trials (Fig. 2).

**Figure 2.**
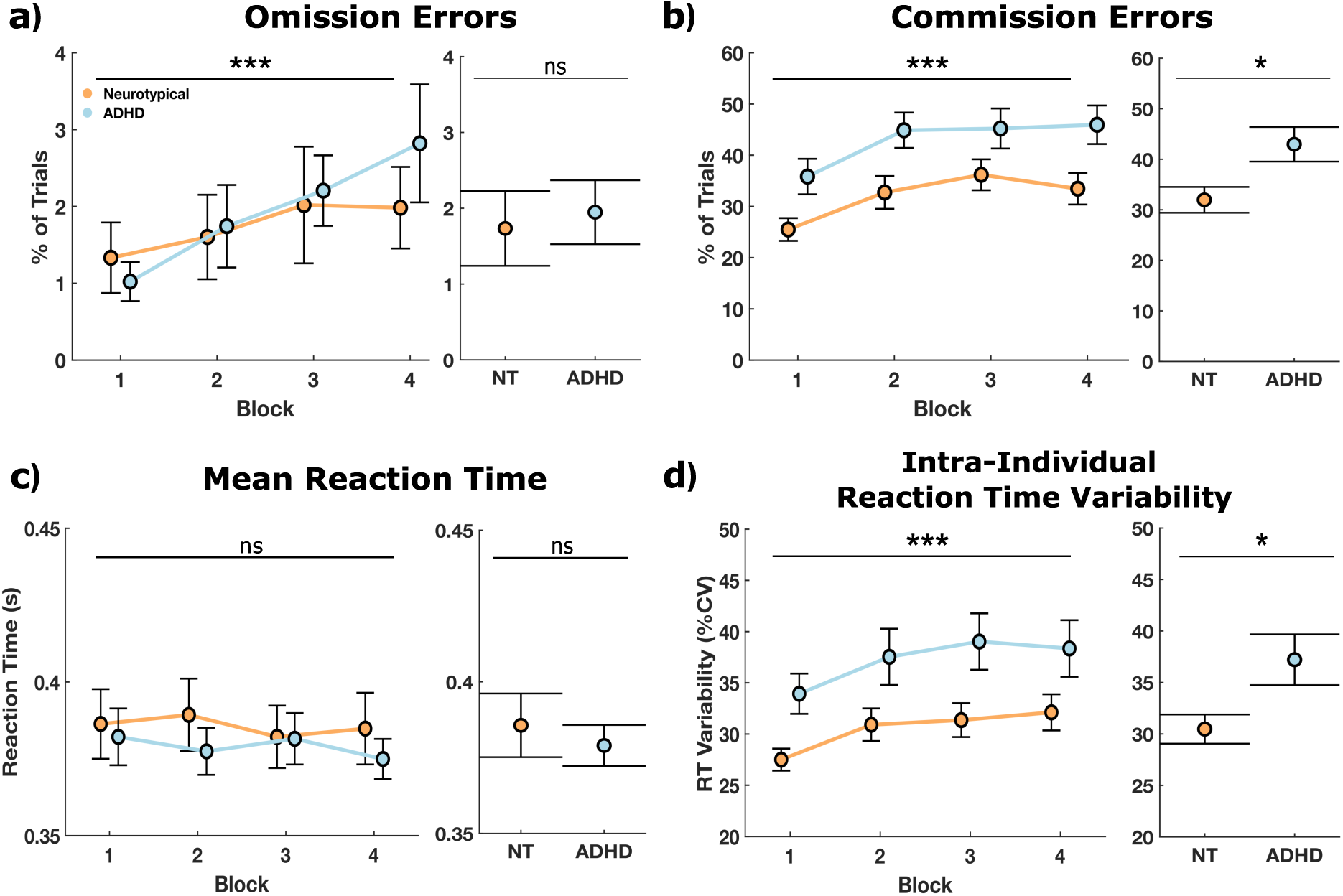
Sustained Attention Response Task (SART) performance results. Performance metrics included: **a.** omission errors, **b.** commission errors, **c.** mean reaction time, and **d.** reaction time variability (intra-individual coefficient of variation of reaction time). The left panels show the mean performance of each group (neurotypical in orange and ADHD in blue) across blocks (N = 4 blocks) with error bars representing the standard error of the performance variables. The right panels show the overall mean and standard error by group. Stars denote the significance level of block effect on performance (left panels) and performance differences between groups (right panels) as determined by FDR-corrected linear mixed-effect models (ns: nonsignificant, \**p* < 0.05, \*\**p* < 0.01, \*\*\**p* < 0.001).

**Figure 3.**
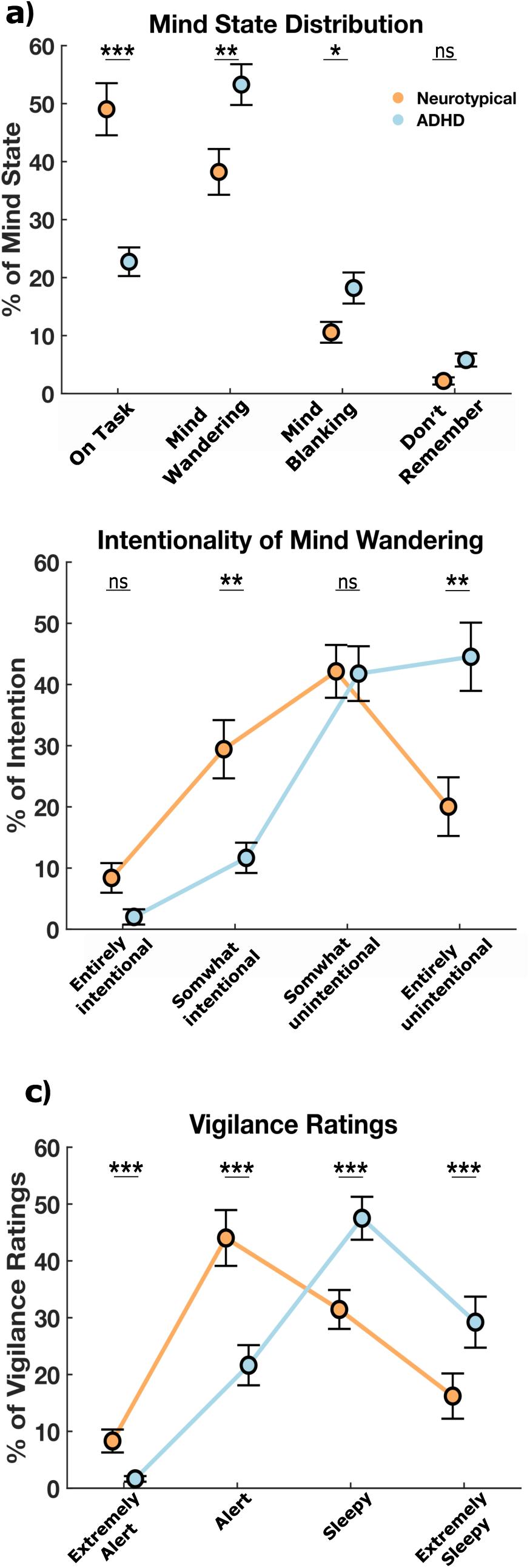
Individuals with ADHD report more MW, less intentionality and higher sleepiness than neurotypical controls (NT). **a.** Distribution of reported mental states across participants and groups. **b.** Distribution of reported intentionality for MW episodes. **c.** Distribution of sleepiness ratings. All panels show mean percentages across probes with error bars representing the standard error of the mean for each group (NT in orange and ADHD in blue). Stars denote the significance level of differences between groups for each category as determined by linear mixed-effect models (ns: nonsignificant, \**p* < .05, \*\**p* < 0.01, \*\*\**p* < 0.001).

**Figure 4.**
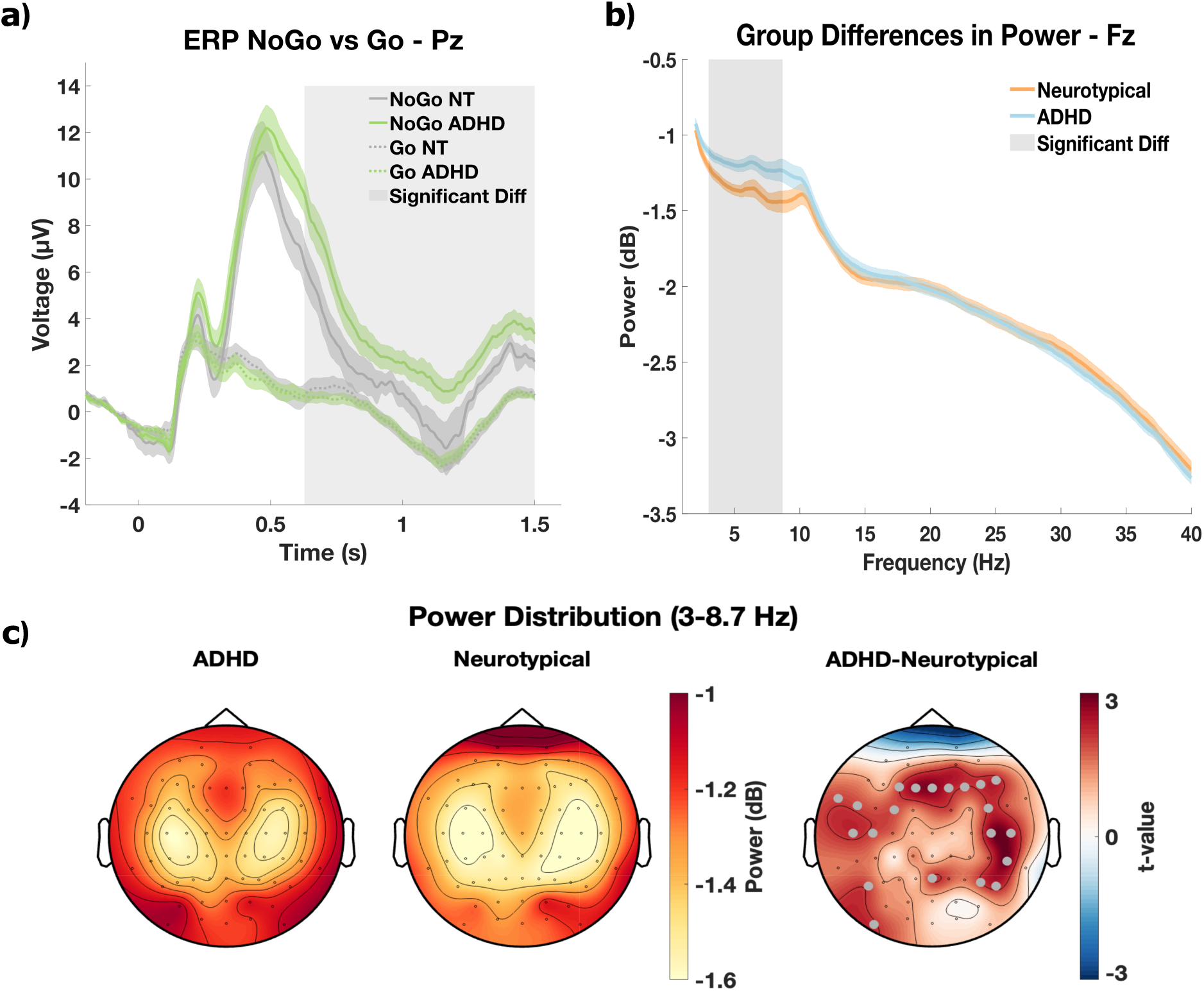
ERPs and power spectrum in ADHD compared with neurotypical controls (NT). **a.** Event-related potential waveforms computed on Pz comparing NoGo (continuous lines) and Go (dashed lines) conditions for NT (grey) and ADHD (green) groups. The shaded areas around each line represent standard error of the mean (SEM). Significant but uncorrected group differences in this NoGo – Go difference were observed between 0.6 and 1.5 s, indicated by the grey shaded region. **b.** Power spectrum density measured at Fz comparing NT (orange) and ADHD (blue) groups across 1–40 Hz. The shaded areas around each line represent SEM. Significant but uncorrected group differences in spectral power were observed between 3 and 8.7 Hz, indicated by the grey shaded region. **c.** Topographical maps of the power distribution across the scalp for the ADHD (left) and NT (middle) groups. The map on the right shows the statistical comparison between the two groups, with *t*-values from unpaired *t*-tests performed at each electrode. Grey dots indicate electrodes with uncorrected *p* < 0.05, highlighting regions where group differences were significant. Note, however, that none of these electrodes survived correction for multiple comparisons (False Discovery Rate).

We also examined the behavioural performance, distribution of mental states and self-report sleepiness ratings according to ADHD presentation (Supp. Fig. 2).

### EEG

#### Data Acquisition and Preprocessing

Participants sat in a dim room, chin on a rest (∼60 cm from a screen). EEG data were recorded using 64 active scalp electrodes with a BrainAmp DC system (Brain Products, Munich, Germany), referenced to FCz, and sampled at 500 Hz. Eye movements were recorded (EyeLink 1000) but not analysed in the current study to focus our analyses on EEG data.

EEG data were preprocessed using a custom MATLAB script focusing on probe-based segmentation. Continuous EEG recordings were segmented from -25 to +2 seconds relative to probe onset, prior to preprocessing. Using the FieldTrip toolbox (Oostenveld et al., 2011), data were demeaned, bandpass filtered (1–40 Hz) with a fourth-order Butterworth filter, notch filtered at 50 and 100 Hz to remove line noise, re-referenced to the average of all channels, and baseline corrected.

We implemented a data-driven approach to automatically identify and remove noisy EEG channels using statistical thresholds based on standard deviation (SD) and kurtosis metrics. For each probe, logarithmic values of SD and kurtosis were computed for each channel. Thresholds for identifying noisy channels were set at three SDs above the mean of non-frontal channels. For frontal channels, thresholds were adjusted using frontal-channel-specific means and standard deviations to account for ocular artefacts. These thresholds identified channels with excessive variability (SD) or abnormal distributions (kurtosis). Channels exceeding these thresholds in more than 50% of probes were identified as noisy (mean number of noisy channels detected per participant ± SEM: 0.22 ± 0.07) and interpolated using the weighted method from FieldTrip with neighbouring channels. Probes were excluded from further analyses if more than 20% of their channels exceeded the statistical thresholds for SD and/or kurtosis (mean number of excluded probes per participant ± SEM: 0.25 ± 0.6).

The preprocessed data were then analysed using Independent Component Analysis (ICA) in EEGLAB for automated classification of artefact components (Delorme & Makeig, 2004; Pion-Tonachini et al., 2019). Before applying the ICA, EEG data were high-passed filtered above 1 Hz to minimise the contamination of slow drifts. Components were automatically removed if they showed a high correlation with eye movements (probability >0.9) or cardiac activity (probability >0.8). The artefact components identified from probe-epoched data were then applied to remove artefacts from trial-epoched data.

Trial-epoched data were segmented around stimulus onset, with epochs from -0.2 to +1.5 seconds after stimulus presentation.

We used probe-epoched data for power spectrum and slow wave analysis, and trial-epoched data for ERP analysis.

#### Event Related Potentials (ERPs)

Trial-epoched data were re-referenced to the average left and right mastoid electrodes (TP9, TP10) and a baseline (-200 to 0 ms) correction was applied. We averaged these time-locked responses by condition to obtain Event-Related Potentials (ERPs) for Go and No-Go trials separately.

#### Spectral analysis

The power spectral density (PSD) was computed for each electrode using the probe-epoched data (20s before each probe). Spectral analysis was performed using FieldTrip’s implementation of the multitaper frequency transform (ft_freqanalysis) with a smoothing frequency of 0.5 Hz across the 1–40 Hz range. Data were padded to the next power of 2 to optimise the Fast Fourier Transform (FFT) computation and was log-transformed (base-10 log).

#### Sleep-like slow waves

Sleep-like slow waves were detected based on established algorithms to detect slow waves in sleep (Riedner et al., 2007) and wakefulness (Andrillon et al., 2021; Pinggal et al., 2022). EEG signals were re-referenced to the average left and right mastoid electrodes (TP9 and TP10) and band-pass filtered between 1–10 Hz using a Chebyshev Type II filter. The start and end of individual waves were defined by two consecutive zero-crossings. For each wave, we extracted a series of parameters including: the most negative and positive peaks and their difference (peak-to-peak amplitude; PTP), duration, and frequency (1/duration). Waves were discarded if they occurred more than 20s before a probe, had a frequency greater than 7 Hz, a positive peak exceeding 75 µV, or occurred within 1s of a high-amplitude event (>150 µV).

High-amplitude slow waves were defined using electrode-specific thresholds derived from the neurotypical group. For each electrode and each neurotypical individual, we computed the 80th percentile of the PTP across all waves. These percentile values were then averaged across the neurotypical group (after excluding participants with values more than 5 standard deviations from the mean). Slow wave thresholds were defined as the average of these individual percentiles for each electrode. Finally, we selected waves exceeding this threshold and classified them as high-amplitude “sleep-like” slow waves. Consistent with previous work (Andrillon et al., 2021; Bernardi et al., 2015; Hung et al., 2013; Quercia et al., 2018; Vyazovskiy et al., 2011), we included waves in the δ-θ range ([1–7] Hz). For each probe, we computed the density (waves/min) of wake slow waves. The threshold selected here (80th percentile) is lower than the 90th percentile used in previous publications focusing on sleep-like slow waves (Andrillon et al., 2021; Pinggal et al., 2022) and was based on the examination of the PTP distributions (Fig. 5a) to avoid being too conservative for the selection of slow waves. We replicated all our analyses with the 90th percentile (Fig. 5 and 6), which provided the same results except for an absence of significant clusters in Fig. 5d.

**Figure 5.**
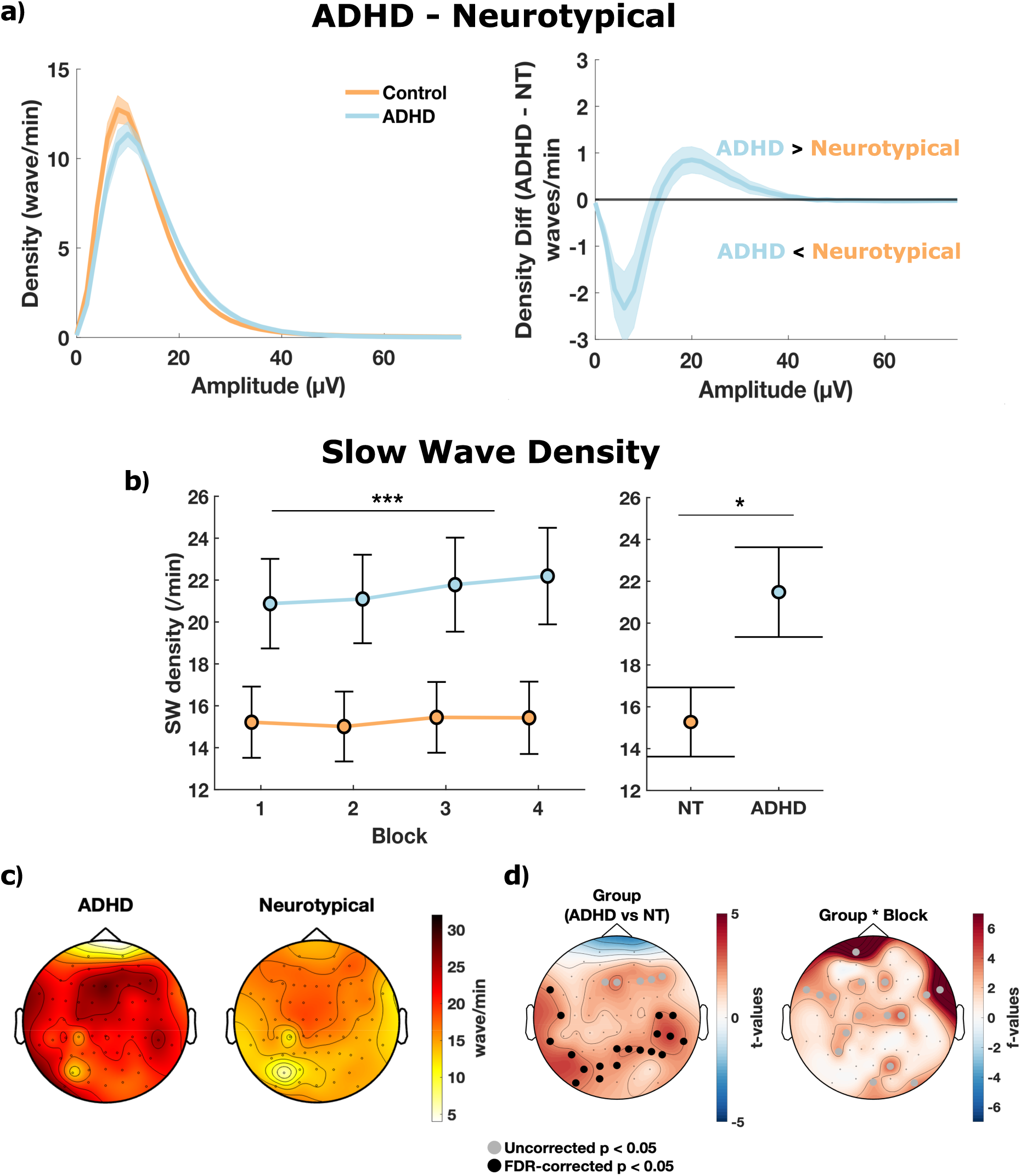
Increased slow wave activity in the ADHD group compared to neurotypical controls (NT). **a.** Left panel: Distribution of slow wave amplitude (waves per minute) showing higher density in NT (orange) but smaller amplitude compared to ADHD (blue) participants. Right panel: Difference wave (ADHD minus NT) with shaded standard error of the mean (SEM) showing amplitude ranges where participants with ADHD exhibited lower or higher amplitudes than NT, highlighting the characteristic shift toward higher amplitude slow waves in ADHD. **b.** The left panel depicts the mean slow wave density of each group (NT in orange and ADHD in blue) across blocks with error bars representing the standard error. The right panel shows the overall mean slow wave density and standard error by group. Stars denote the significance level of block effect on slow wave density (left panels) and differences between groups (right panels) as determined by linear mixed-effect models (ns: nonsignificant, \**p* < 0.05, \*\**p* < 0.01, \*\*\**p* < 0.001). **c.** Topographical maps of the slow wave density distribution across the scalp for the ADHD and NT groups. **d.** Statistical comparison maps showing the *t*-values resulting from a linear mixed effect model performed for each electrode for group main effect (ADHD vs NT, left) and group by block interaction (right). Grey dots indicate significant electrodes with uncorrected *p* < .05, while black dots represent electrodes with significant differences after cluster permutation (1000 permutations, cluster α = .05).

**Figure 6.**
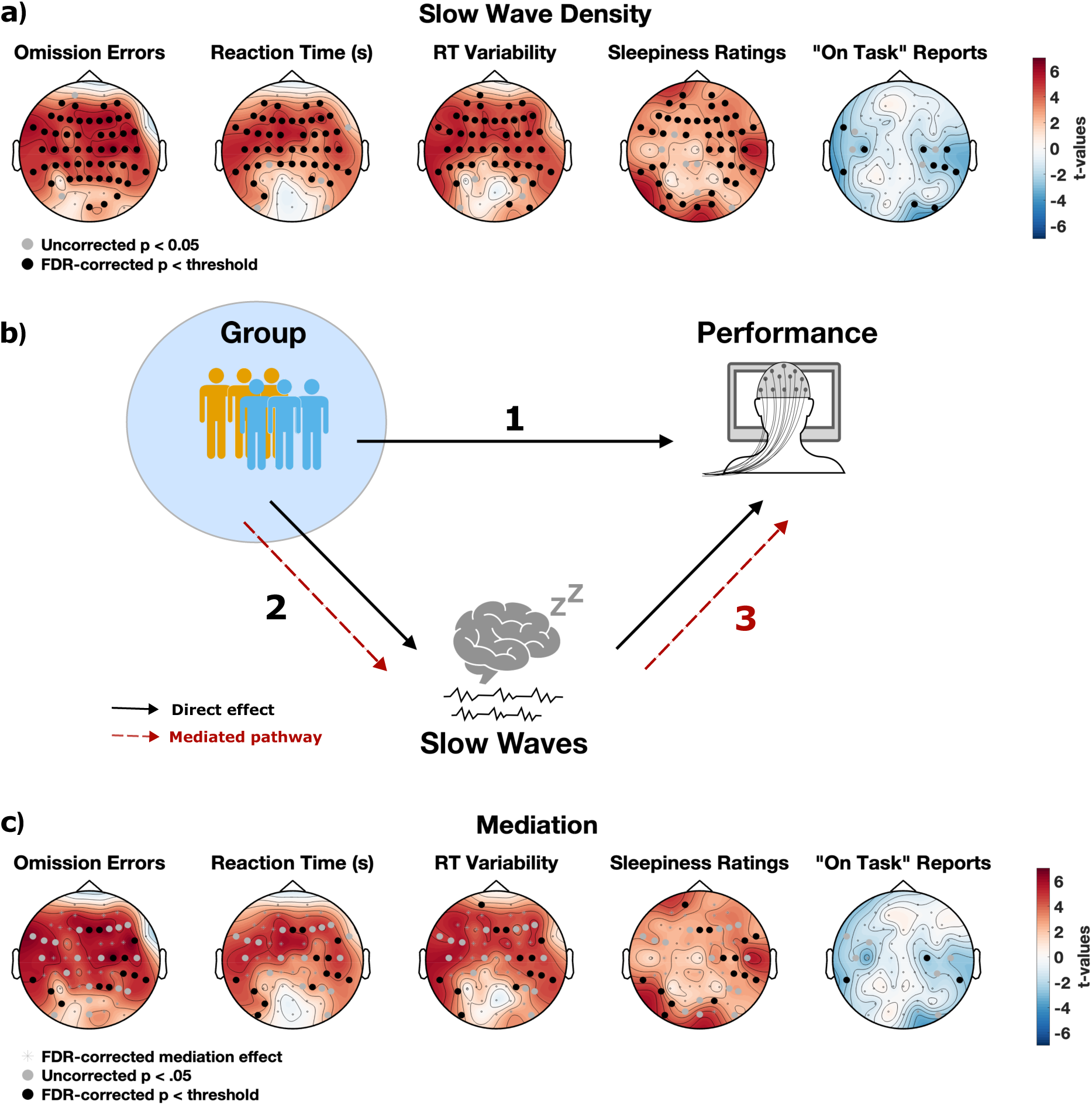
Slow wave density correlates with and mediates task performance. **a.** Topographical maps showing *t*-values from linear mixed effect models testing the association between slow wave density (SWD) and behavioural measures: omission errors, reaction time, reaction time variability (ICVRT), subjective sleepiness ratings, and on-task reports. Positive *t*-values (red) indicate positive associations, while negative *t*-values (blue) indicate negative associations. **b.** Conceptual diagram of the mediation analysis examining how slow waves mediate the relationship between group status (neurotypicals in orange, ADHD in blue) and task performance. **c.** Topographical maps showing *t*-values of the mediation effect for each performance measure. Significant electrodes are marked as black, bold dots (FDR-corrected *p* < threshold) or grey (uncorrected *p* < .05). FDR correction was applied jointly across analyses in Figures 6a and 6c.

### Statistical analyses

To compare demographics, we conducted *t*-tests (age, education, CAARS and ESS) and a chi-square test (sex; *p-*values in Table 1).

Linear mixed effect models (LMEs) in MATLAB R2022b (‘fitlme’, Statistics and Machine Learning Toolbox) assessed group effects (categorical fixed effect) on commission errors, omission errors, RTs (correct Go trials), RT variability, *d*-prime, mental states, sleepiness ratings and slow wave density. Errors and RTs were computed across trials, RT variability, *d*-prime and mental states across blocks, and sleepiness ratings and slow wave density across probes. Models included subject identity as a random effect, and block and group as fixed effects, with some including a block-by-group interaction. Model comparisons were conducted using likelihood ratio tests to determine the best-fitting model. To estimate the effect of categorical variables, we ran an ANOVA on the model (‘anova’ function) and reported the F-values along with the associated degrees of freedom and *p*-value. We also report the models’ coefficients (β) and the associated standard errors (SE) to estimate the strength and direction of the effect of a specific variable. For performance and mental state analyses, *p*-values were adjusted using False Discovery Rate (FDR) control via the Benjamini Hochberg method (α = .05).

Chi-squared tests assessed group differences across sleepiness levels (1–4), and the association between sleepiness and mental state reports. For the latter and for each mental state / sleepiness pair, standardised residuals were inspected post hoc and values greater than ±1.96 were considered as indicating significant over- or under-representation.

To estimate the effect size of the association between sleepiness and mental states (two categorical variables), we computed Cramer’s *V*. To estimate the effect size of the group on each sleepiness rating or mental state (post-hoc test), we used the Phi coefficient. Independent *t*-tests (FDR Benjamini-Hochberg-corrected) examined the relationship between intentionality and MW, with corresponding Cohen’s *d* effect sizes reported. Exploratory analyses assessed whether performance and mental state reports differed by ADHD presentation (Supp. Fig. 2).

Examining ERPs in the ADHD and neurotypical groups revealed a clear P300 component (400ms post-stimulus; Figure 4) for NoGo trials compared to Go trials. To quantify the group differences, we focused on representative electrodes (Pz and Fz). For spectral analysis, we computed PSD across the 1–40 Hz range. We used ANOVAs to assess group differences over a specific time (ERPs) or frequency (PSD) window. A cluster-based permutation approach (Maris & Oostenveld, 2007) was used to correct for multiple comparisons (1000 permutations, cluster α = .05, Monte-Carlo α = .05).

For slow wave density, we fitted LMEs with group, block and their interaction as fixed effects, and participant ID as a random effect. A cluster-based permutation approach (Maris & Oostenveld, 2007) was used to correct for multiple comparisons (1000 permutations, cluster α = .05, Monte-Carlo α = .05).

We conducted a form mediation analysis to test whether the relationship between group and task performance could be explained, at least in part, by individual differences in sleep-like slow wave activity. To assess whether performance was associated with slow wave density independently of group, we examined their relationship after statistically controlling for group (ADHD vs. neurotypical). Establishing that (1) group modulates performance, (2) group modulates slow wave density, and (3) slow wave density and performance are still correlated after statistically controlling for the influence of group, can be interpreted as evidence for a mediation effect of group on performance by slow waves. In practice, we first fit separate LMEs for performance and slow wave density at each electrode, including group, block and their interaction as fixed effects and subject identity as a random intercept. Residuals from each model were extracted, representing variance unexplained by group and block. We then computed correlations between the residuals of these models to determine whether performance variance was associated with slow wave density after regressing out the effect of group. We restricted our mediation analyses to performance variables that showed a significant group effect (omission errors, RT, RT variability, sleepiness ratings and on-task reports). For each of these variables and each electrode, evidence for mediation of the group effect on performance via slow wave density was defined as (1) a significant effect of group on slow

wave density, and (2) a significant correlation between the residuals of slow wave density and the residual of the performance variable. To control for multiple comparisons across electrodes and variables, FDR correction was applied: *p*-values from tests examining slow wave effects on performance and from mediation analyses were aggregated for FDR correction.

## Results

### Behaviour

The best fitting models for commission errors, RT, *d*-prime and RT variability included group and block number as fixed effects, and subject identity as a random effect. For omission errors, the best fitting model included a group × block interaction. FDR correction was applied using the Benjamini-Hochberg procedure (α = .05).

Although there was no overall group effect for omission errors (F(1, 118,740) = 0.81, *p* = .434), omission errors increased across blocks (F(1, 118,740) = 25.50, *p* < .001) and there was a significant group × block interaction (F(1, 118,740) = 21.91, *p* < .001; see Table 2 for details). This interaction was driven by a steeper increase in omission errors across blocks in the ADHD group (β = 0.0057, SE = 0.0005 per block) compared with the neurotypical group (β = 0.0025, SE = 0.0007 per block) (Fig. 2). Commission errors were higher in the ADHD than the neurotypical group (F(1, 14,830) = 6.98, *p* = .018) and increased over time across groups (F(1, 14,830) = 78.82, *p* < .001), although there was no significant group × block interaction (χ²(1) = 0.30, *p* = .586).

**Table 2.** LME fixed effects statistics for SART performance.

| Performance | | Estimate<br>($\beta$) | SE | *t*-value | df | uncorrected<br>*p*-Value | FDR-corrected<br>*p*-Value |
| --- | --- | --- | --- | --- | --- | --- | --- |
| **Omission errors** | Intercept | 0.011 | 0.0046 | 2.40 | 118,740 | 0.016 |  |
|  | Group (ADHD) | -0.0058 | 0.0065 | -0.90 | 118,740 | 0.368 | 0.434 |
|  | **Block** | **0.0032** | **0.00069** | **5.05** | **118,740** | **< .001** | **< .001** |
|  | **Group × Block** | **0.0032** | **0.00069** | **4.68** | **118,740** | **< .001** | **< .001** |
| **Commission errors** | Intercept | 0.25 | 0.031 | 8.01 | 14,830 | < .001 |  |
|  | **Group (ADHD)** | **0.11** | **0.042** | **2.64** | **14,830** | **.008** | **0.018** |
|  | **Block** | **0.03** | **0.0033** | **8.88** | **14,830** | **< .001** | **< .001** |
| ***d*-prime** | Intercept | 2.61 | 0.17 | 14.94 | 249 | < .001 |  |
|  | **Group (ADHD)** | **-0.47** | **0.22** | **-2.13** | **249** | **.034** | **.049** |
|  | **Block** | **-0.15** | **0.030** | **-5.18** | **249** | **< .001** | **< .001** |
| **criterion** | Intercept | -0.95 | 0.05 | -20.40 | 249 | < .001 |  |
|  | **Group (ADHD)** | **-0.12** | **0.05** | **-2.36** | **249** | **.019** | **.031** |
|  | Block | -0.004 | 0.01 | -0.38 | 249 | .70 | .763 |
| **Reaction Time** | Intercept | 0.39 | 0.0088 | 44.16 | 112,970 | < .001 |  |
|  | Group (ADHD) | 0.0005 | 0.012 | 0.04 | 112,970 | .968 | .968 |
|  | Block | 0.0006 | 0.00034 | 1.73 | 112,970 | .084 | .109 |
| **Reaction Time**<br>**Variability**<br>**(ICVRT)** | Intercept | 0.10 | 0.0083 | 12.59 | 249 | < .001 |  |
|  | **Group**<br>**(ADHD)** | **0.067** | **0.028** | **2.43** | **249** | **.016** | **.029** |
|  | **Block** | **0.015** | **0.0025** | **5.70** | **249** | **< .001** | **< .001** |

*Note.* **Bold** = significant main effect after $p$ -values were FDR-corrected using the Benjamini-Hochberg method. ADHD = Attention Deficit Hyperactivity Disorder; ICVRT = Intra-Individual Coefficient of Variation of Reaction Time.

Signal detection analyses revealed a significantly lower *d*-prime in the ADHD group compared to the neurotypical group (F(1, 249) = 4.54, p = .049). Additionally, the ADHD group had a lower response criterion compared to the neurotypical group (F(1, 249) = 5.56, *p* = .031), reflecting a more liberal response strategy in ADHD and a relatively more conservative strategy in the neurotypical group.

RT analyses showed no significant group (F(1, 112,970) = 0.0016, *p* = .968) or block effects (F(1, 112,970) = 2.99, *p* = .109), indicating stable response speeds over time. However, RT variability (ICVRT) was higher in the ADHD group compared to the neurotypical group (F(1, 249) = 5.90, *p* = .029). Additionally, RT variability increased over time regardless of group (F(1, 249) = 32.50, *p* < .001), suggesting greater overall response inconsistency, with ADHD participants exhibiting higher variability throughout.

### Mental states

We examined mental state reports (on task, MW, MB and “*don’t remember”*) using block-level LMEs (Fig. 3). Likelihood ratio tests indicated that the best-fitting model included group and block as fixed effects and subject identity as a random effect. For "*don’t remember*" reports, the best-fitting model included a group × block interaction. FDR correction was applied using the Benjamini-Hochberg procedure (α = .05).

There was a significant main effect of group on on-task reports (F(1, 249) = 28.56, *p* < .001), with the ADHD group reporting fewer on-task states than the neurotypical group (see Table 3 for details). Additionally, in both groups, on-task reports decreased over time (F(1, 249) = 37.81, *p* < .001). The significant main effect of group on MW (F(1, 249) = 8.68, *p* = .004) and MB (F(1, 249) = 5.91, *p* = .016) were driven by higher reports in the ADHD group than the neurotypical group. There was a significant main effect of block on MW (F(1, 249) = 27.69, *p* < .001), but not MB (F(1, 249) = 1.10, *p* = .295). "*Don’t remember*" responses had non-significant group (F(1, 248) = 0.03, *p* = .864) and block effects (F(1, 248) = 0.22, *p* = .642). However, the group × block interaction was significant (F(1, 248) = 5.44, *p* = .02), indicating that *“Don’t remember”* reports increased more across blocks in the ADHD group compared to the neurotypical group

**Table 3.** LME fixed effects statistics for mental state reports.

| mental states | | Estimate<br>($\beta$) | SE | *t*-value | df | uncorrected<br>*p*-Value | FDR-corrected<br>*p*-Value |
| --- | --- | --- | --- | --- | --- | --- | --- |
| **On Task** | Intercept | 0.63 | 0.042 | 15.09 | 249 | < .001 |  |
|  | **Group**<br>**(ADHD)** | **-0.26** | **0.049** | **-5.34** | **249** | **< .001** | **< .001** |
|  | **Block** | **-0.055** | **0.009** | **-6.15** | **249** | **< .001** | **< .001** |
| **Mind**<br>**Wandering** | Intercept | 0.25 | 0.044 | 5.67 | 249 | < .001 |  |
|  | **Group**<br>**(ADHD)** | **0.15** | **0.051** | **2.95** | **249** | **.004** | **.008** |
|  | **Block** | **0.052** | **0.010** | **5.26** | **249** | **< .001** | **< .001** |
| **Mind**<br>**Blanking** | Intercept | 0.125 | 0.029 | 4.28 | 249 | < .001 |  |
|  | **Group**<br>**(ADHD)** | **0.076** | **0.03** | **2.43** | **249** | **.016** | **.028** |
|  | Block | -0.0079 | 0.0076 | -1.05 | 249 | .295 | .379 |
| **“Don’t**<br>**Remember”** | Intercept | 0.016 | 0.015 | 1.07 | 248 | .285 |  |
|  | Group<br>(ADHD) | -0.0036 | 0.021 | -0.17 | 248 | .864 | .864 |
|  | Block | 0.0023 | 0.0049 | 0.47 | 248 | .642 | .722 |
|  | **Group ×**<br>**Block** | **0.016** | **0.0068** | **2.33** | **248** | **.021** | **.031** |

*Note.* **Bold** = significant main effect after *p*-values were FDR-corrected using the Benjamini-Hochberg method.

For MW intentionality (independent *t*-tests, FDR Benjamini-Hochberg correction α = .05), neurotypical controls reported more instances of "*somewhat intentional*" MW (*t*(60) = 3.43, *p* = .006, Cohen’s *d* = 0.86), whereas participants with ADHD reported more *"entirely unintentional*" episodes (*t*(60) = -3.37, *p* = .006, Cohen’s *d* = -0.86; see Fig. 3). The trend for group differences in "*entirely intentional*" (*t*(60) = 2.42, *p* = .05, Cohen’s *d* = 0.61) did not survive correction. No significant group differences were observed for “*somewhat unintentional*” MW (*t*(60) = 0.06, *p* = 1.98, Cohen’s *d* = 0.02).

### Self-Reported Sleepiness Ratings

To assess self-reported sleepiness ratings, the best fitting LME model, as determined from likelihood ratio tests, included group and block as fixed effects and subject identity as a random effect. There was a significant main effect of group (F(1, 2517) = 17.12, *p* < .001) with participants with ADHD reporting higher sleepiness overall compared to neurotypical individuals (see Table 4 for detailed LME results). There was also a significant main effect of block (F(1, 2517) = 288.05, *p* < .001) with sleepiness increasing over time.

**Table 4.** LME fixed effects statistics for self-reported sleepiness ratings.

| | | Estimate<br>($\beta$) | SE | *t*-value | df | uncorrected<br>*p*-Value | FDR-corrected<br>*p*-Value |
| --- | --- | --- | --- | --- | --- | --- | --- |
| **Sleepiness Ratings** | Intercept | 2.08 | 0.09 | 23.47 | 2517 | < .001 |  |
|  | **Group (ADHD)** | **0.49** | **0.12** | **4.14** | **2517** | **< .001** | **< .001** |
|  | **Block** | **0.19** | **0.01** | **16.97** | **2517** | **< .001** | **< .001** |

*Note.* **Bold** = significant main effect after *p*-values were FDR-corrected using the Benjamini-Hochberg method.

Post-hoc chi-square tests revealed significant group differences across all sleepiness levels (Fig. 3). Neurotypical individuals reported higher proportions of alert ratings, particularly at level 1 (extremely alert; χ²(1, 2520) = 59.82, *p* < .001, φ = 0.15), and level 2 (alert; χ²(1, 2520) = 143.59, *p* < .001, φ = 0.24). Conversely, the ADHD group reported more sleepiness ratings with significant group differences at level 3 (sleepy; χ²(1, 2520) = 67.82, *p* < .001, φ = 0.16) and level 4 (extremely sleepy; χ²(1, 2520) = 60.53, *p* < .001, φ = 0.15). All effects remained significant after FDR correction (α = .05). Effect sizes indicate small-to-moderate associations between group and sleepiness ratings across levels (φ = 0.15–0.24).

To assess the association between sleepiness ratings and mental state reports, a chi-squared test of independence was conducted. A significant association was found, χ²(9, 2560) = 569.83, *p* < .001, with a moderate effect size (Cramer’s *V* = 0.27). Post hoc analysis of standardised residuals identified specific cell-level contributions: on-task reports were significantly over-represented when participants reported being extremely alert and alert, and under-represented when extremely sleepy and sleepy, whereas mind wandering and mind blanking were over-represented at higher sleepiness levels. Table 5 presents the percentage distribution of mental states across sleepiness ratings.

**Table 5.** Percentage distribution of mental state reports by self-reported sleepiness ratings.

|  | *N* | On Task | Mind<br>Wandering | Mind<br>Blanking | Don't<br>Remember |
| --- | --- | --- | --- | --- | --- |
| Extremely Alert | 155 | **90.97%** | **7.74%** | **1.29%** | **0%** |
| Alert | 832 | **54.81%** | **38.46%** | **4.93%** | **1.80%** |
| Sleepy | 998 | **28.46%** | 48.40% | **19.34%** | 3.81% |
| Extremely Sleepy | 575 | **10.09%** | **59.30%** | **22.26%** | **8.35%** |

*Note.* *N* indicates the number of probes within each sleepiness rating. Bolded percentages indicate statistically significant over- or under-representation (standardised residuals $\geq \pm 1.96$ ) based on post-hoc chi-squared residual analysis. Percentages reflect the distribution of mental state reports within each sleepiness category.

### ERP and Power Spectrum

As in previous studies using the SART, we observed that NoGo trials elicited a strong P300 component compared to Go trials. We thus examined group differences in ERPs for NoGo and Go trials at Pz using ANOVAs for each time point. Uncorrected tests showed differences in the 0.63–1.5 s window (Fig. 4a), on a time window following the P300 (see Discussion). However, these differences did not survive a cluster-based permutation correction (1,000 permutations, α = .05).

For spectral analysis, we compared PSD (1–40 Hz) at Fz using the same approach. While no group differences survived correction, uncorrected results indicated differences in the 3–8.7 Hz range (Fig. 4b). Exploratory analyses across all electrodes (Fig. 4c) identified significant clusters primarily in frontal and temporal regions, uncorrected for multiple comparisons.

### Slow waves in wakefulness

Prior to thresholding the waves, the distribution of all the waves detected revealed that the ADHD group had fewer low-amplitude slow waves but more higher-amplitude waves compared to neurotypicals (Fig. 5a). This shift between the two group is observed at moderate amplitude (∼15µV), which prompted us to avoid being too conservative when selecting slow waves based on their amplitude (see Methods). We then selected the highest amplitude slow waves, based on amplitude thresholds computed at the electrode and subject level for the neurotypical group and averaged across subjects (see Methods). The global SW density (averaged across electrodes) indicate a sustained and significant increase in the ADHD group (Fig. 5b). Model comparisons indicated a significant interaction between group and block.

Topographies illustrate an increase in slow wave density across the scalp in the ADHD group compared to the neurotypical group (Fig. 5c). We then examined group differences in slow wave density by fitting LMEs at each electrode. To control for multiple comparisons, we completed a cluster permutation (1000 permutations, cluster α = .05; see Methods for further details). Slow wave density was modelled as the dependent variable, with group, block, and their interaction as fixed effects, and subject identity as a random effect. Results are reported as topographies in Figure 5d and revealed higher slow wave density in the ADHD group, particularly in parieto-temporal electrodes compared to neurotypicals. The significant electrodes found for the group-by-block interaction did not survive the cluster permutation analysis.

To investigate how slow wave activity in wakefulness relates to SART performance, LMEs were implemented. Dependent variables included performance measures (commission errors, omission errors, mean RT and RT variability) and probe responses (MW, on-task reports, and sleepiness ratings). Fixed effects included block, slow wave density, and their interactions with electrode location, with subject identity as a random effect.

Results are reported as topographies in Figure 6a which showed that higher global slow wave density was associated with more omission errors, slower mean RTs, and greater RT variability, while commission errors showed no relationship. Increased slow wave density in frontal and occipital regions were also predictive of higher sleepiness ratings (Fig. 6a). While MW showed no association with slow wave density, lower slow wave density in temporal regions was associated with more on-task reports.

#### Mediation analysis

We conducted electrode-wise mediation analyses to examine if slow wave density in wake mediated the relationship between ADHD and task performance, controlling for block effects. As detailed in the Methods section, the mediation of the effect of group on performance by the density of slow waves in electrode *k* is tested by verifying that there is a (1) a significant effect of group on behaviour, (2) a significant effect of group on slow wave density for electrode *k*, and (3) a significant correlation between the slow wave density and performance after removing the variance explained by the group effect (correlation between residuals; see Fig. 6). This is to show that slow wave density and performance are associated even after the removal of the group effect, suggesting that the correlation between the two variables is not simply driven by the group effect. Significant mediation effects (Fig. 6c) indicated that group differences in slow wave density mediated the link between ADHD and performance, particularly for omission errors, mean RT, and RT variability, with effects most prominent in frontal and centro-parietal regions (FDR-corrected *p* < threshold). Sleepiness ratings also showed significant mediation in right centro-parietal electrodes, while on-task reports showed minimal mediation, with few electrodes reaching significance.

## Discussion

### Attentional Difficulties in ADHD: Behavioural and Cognitive Manifestations

The SART findings highlight the cognitive challenges experienced by adults with ADHD, revealing significantly higher commission errors, in line with the condition’s elevated impulsivity. Beyond impulsivity, increased RT variability across blocks in the ADHD group suggests worsening attentional control over time – a marker of attentional dysregulation commonly used in ADHD research (Kofler et al., 2013). These findings support and extend previous research (Bozhilova et al., 2020; Madiouni et al., 2020; Van Den Driessche et al., 2017), providing insights into potential time-on-task effects that reflect increased fatigue susceptibility and possible trait-like hypo-arousal in ADHD.

Performance across task blocks revealed a decline in cognitive control (Fig. 2). Irrespective of group, commission errors increased across blocks, indicating a decline in top down control over time, which here manifests as a failure of sustained attention to the task (Gunzelmann et al., 2011; Tucha et al., 2017). While omission errors did not significantly differ between groups, both groups exhibited an increase over time, with the ADHD group showing a steeper increase (significant interaction between group and block, see Results), consistent with an increased cognitive fatigability in ADHD. Despite these differences, both groups maintained comparable mean RTs (Bozhilova et al., 2020; Madiouni et al., 2020), though the ADHD group showed more variability in response times.

Beyond objective performance metrics, thought probe analyses revealed distinct subjective experiences between groups (Fig. 3). Neurotypical individuals reported more on-task focus, while those with ADHD reported more MW (Madiouni et al., 2020; Seli et al., 2015). This difference in subjective experience extends beyond frequency, unveiling nuanced distinctions in attentional disengagement. Participants with ADHD primarily experienced more “*entirely unintentional”* MW whereas neurotypical participants reported more *“somewhat intentional”* episodes, suggesting that there is indeed a difference in neural mechanisms governing attentional control (Seli et al., 2015). Moreover, MB reports in the ADHD group were also higher, and thus may reflect underlying executive disengagement, consistent with previous studies reporting elevated MB in this population (Madiouni et al., 2020; Van Den Driessche et al., 2017). In line with this, a significant interaction was found between group and block for *“Don’t remember”* reports, with individuals with ADHD showing a gradual accumulation of these responses across time, indicative of declining meta-awareness or monitoring capacity as the task unfolds.

These results indicate altered metacognitive regulation of attentional states in ADHD. This aligns with findings by Franklin et al. (2017), who noted that neurotypical individuals with low ADHD symptoms strategically allocate attentional resources. This strategic cost-benefit assessment was notably absent in individuals with high ADHD symptoms, potentially reflecting disruptions in higher-order executive functions necessary for regulating attentional shifts. Consequently, the greater frequency and lower intentionality of MW in ADHD may reflect difficulties in executive functions required for regulating attentional shifts.

Across all participants, mental states also varied with subjective sleepiness. On-task reports were more frequent when participants felt alert, while MW and MB increased as sleepiness levels rose. Although nearly half of ‘sleepy’ probes were associated with MW (48.4%), this was consistent with expected levels, suggesting it reflects a typical level of attentional disengagement under moderate fatigue. In contrast, extremely sleepy ratings indicated a disproportionate increase in off-task states, suggesting that increasing sleepiness may further compromise the ability to maintain focused attention.

Additionally, task-based sleepiness ratings revealed that participants with ADHD reported higher sleepiness and fewer alert ratings compared to neurotypical participants. Interestingly, this contrasts with the non-significant group differences on the Epworth Sleepiness Scale. This pattern suggests that while subjective sleepiness fluctuates more dynamically in ADHD, consistent with Helfer et al. (2020) who observed elevated daytime sleepiness in adults with ADHD during a SART, global, trait-level sleepiness did not significantly differ between the two groups. Together, these results highlight that sustained attentional efforts in ADHD result in increased behavioural variability and greater fluctuations in vigilance over time.

### Sleep-like slow wave activity in wakefulness: a neurophysiological mechanism of attentional difficulties in ADHD?

Our EEG analyses revealed distinct slow wave activity patterns in individuals with ADHD, characterised by higher amplitude and density in parieto-temporal electrodes (Fig. 5). Emerging research suggests that sleep-like slow waves can intrude into specific brain regions during wakefulness (Andrillon et al., 2021; Pinggal et al., 2022; Vyazovskiy et al., 2011), and our findings support the hypothesis of “sleepier brains” in ADHD (Andrillon et al., 2019). This indicates that slow wave activity during wake occurs more frequently in ADHD, offering a potential neurophysiological explanation for attentional challenges.

To further explore the effects of sleep-like slow wave activity in wake, we examined its impact on sustained attention. We observed a global increase in slow wave density correlated with several performance decrements: higher omission errors, slower RT, and greater RT variability (Fig. 6). The effect was particularly evident in central electrodes where increased slow wave activity appeared to impair response preparation and execution. This pattern corroborates previous research demonstrating how slow wave activity during wakefulness disrupts cognitive processing (Andrillon et al., 2021; Pinggal et al., 2022). However, we did not find a correlation between SW and commission errors in this dataset. Yet, the stronger presence of this neurophysiological marker in individuals with ADHD provides evidence that it may underlie some of their characteristic attentional difficulties. Furthermore, the global increase in slow wave density was associated with elevated subjective sleepiness ratings, supporting the notion that slow waves reflect accumulating sleep pressure, affecting both objective performance and subjective experience. Lastly, a negative correlation between local sleep and subjective attentional focus was observed: elevated local sleep corresponded with fewer “*on-task*” reports. This inverse relationship highlights that increased slow wave density not only impacts objective performance, but also subjectively diminishes task engagement.

#### Slow-wave activity as a mediating factor

We further explored whether slow wave density mediated the relationship between ADHD diagnosis and SART performance through electrode-wide mediation analysis. The strongest mediation effects were observed for omission errors, RT, and RT variability, with significant electrodes primarily clustered in frontal and centro-parietal regions (Fig. 6c). These findings indicate that slow wave activity mediates the link between ADHD and behaviour, with ADHD influencing performance through specific alterations in slow waves. While mediation effects for subjective “*on-task*” reports were weaker, the consistency of results across multiple objective measures – from RT variability, omission errors to overall RT and sleepiness ratings – suggests slow wave activity may serve as a neurophysiological bridge between ADHD and its behavioural manifestations.

Crucially, this also highlights sleep-like slow waves as a promising target for intervention. Pharmacological treatments commonly used in ADHD, such as methylphenidate and atomoxetine, act on dopaminergic and noradrenergic systems that regulate arousal (Bymaster, 2002; Chamberlain et al., 2009; Koda et al., 2010; Kowalczyk et al., 2019). These medications not only improve performance, but also reduce the occurrence of sleep-like slow waves, particularly over frontal and central scalp regions (Pinggal et al., 2022). By identifying local sleep as a mediator, these findings advance our understanding of the neural mechanisms of ADHD and highlight a potential biomarker that could guide interventions targeting sleep-like slow waves in specific neural regions. In addition to pharmacological approaches, addressing sleep difficulties in ADHD, such as through behavioural interventions like Cognitive Behavioural Therapy for Insomnia (CBT-I), may aid in reducing the occurrence of sleep-like slow waves during wakefulness. Improving sleep quality and regulation, could, in turn, support daytime arousal and attentional stability.

#### Disentangling trait and state arousal impairments in ADHD

The observed hypo-arousal effects in ADHD raise important questions about their underlying mechanisms. These effects may stem from a chronic state of reduced arousal, as indicated by greater slow wave activity, and increased self-reports of sleepiness during the task in the ADHD group. These effects could also, or instead, arise from a use-dependent mechanism, with individuals with ADHD experiencing a faster onset of fatigue during sustained effort. This view is supported by a significant group × time interaction in our analyses, whereby omission errors increased more steeply across blocks in the ADHD group. These two accounts are not mutually exclusive, and future research should endeavour to disentangle trait-like versus task-induced components of hypo-arousal in ADHD.

#### Supplementary neural signatures: ERP and Spectral perspectives

Exploratory analyses of ERPs and PSD provided additional neural insights. We did not observe a reduction of the P300 component in contrast with previous studies (Szuromi et al., 2011). However, individuals with ADHD showed an enhanced Late Positive Potential (LPP) (Jaswal et al., 2019), which could reflect enhanced post-perceptual or error-related cognitive processes in ADHD individuals. However, these differences did not reach statistical significance after correcting for multiple comparisons. Similarly, spectral analyses at Fz revealed no significant group differences after cluster-based correction but uncorrected statistics show increased power in the delta/theta range, consistent with our findings on slow waves. This absence of a significant power effect may be explained by the differential modulation of slow waves in ADHD: while high-amplitude slow waves are increased, low-amplitude slow waves are reduced (Figure 5a). Consequently, a conventional power analysis, which aggregates both high- and low-amplitude slow waves, may obscure these opposing effects, leading to only a modest overall increase in power for the ADHD group. This suggests that focusing on sleep-like slow waves may serve as a more sensitive and specific neural marker for distinguishing ADHD from neurotypical individuals.

### Limitations and future directions

While our findings provide insights into the neurophysiological mechanisms underlying attentional challenges in ADHD, several limitations must be acknowledged. Despite a 72-hour washout period, there is limited research on the lingering effects of stimulants on cognitive performance. Additionally, our predominantly female cohort contrasts with typical ADHD research cohorts which tend to favour males. Nevertheless, by focusing on this underrepresented group, our study addresses critical research gaps in the literature regarding how ADHD manifests in females. Females with ADHD often present with more inattentive symptoms and may employ better masking strategies than males, while those who are hyperactive tend to exhibit verbal rather than physical overactivity (Faraone et al., 2024; Hinshaw et al., 2022). Our study may offer insights into female-specific characteristics of local sleep-like activity and attentional fluctuations in ADHD. Future studies should aim for a more gender-balanced cohort to determine generalisability and potential gender differences in local sleep manifestations. Moreover, the sleep behaviour of participants were not measured in the days leading up to their lab visit. Thus, we cannot exclude the influence of habitual, or at least recent, sleep patterns from our findings.

Lastly, as a cross-sectional study, we cannot determine whether local sleep represents a stable trait marker (i.e., increased slow wave activity in wakefulness as a neurophysiological characteristic of ADHD) or a transient state induced by the task. Longitudinal research and diverse cognitive paradigms (e.g., working memory or selective attention tasks) could help clarify the stability and specificity of these neural patterns. Since sleep and vigilance are not only under the control of time spent awake (homeostatic pressure) but also of the position within the circadian cycle (Borbély, 2022), future studies should control for the timing of tasks and circadian profiles. Finally, given the heterogeneity within ADHD presentations, future research can also aim to investigate whether slow wave activity during wakefulness differs between each of the three presentations.

## Conclusion

This study offers a comprehensive investigation into the neurophysiological underpinnings of sustained attention difficulties in ADHD, with a novel focus on local sleep-like activity during wakefulness. Participants with ADHD demonstrated greater attentional lapses, evidenced by task performance and MW reports. Critically, participants with ADHD exhibited higher slow wave density, particularly in parieto-temporal electrodes. Mediation analyses revealed that these slow waves not only predict, but also mediate the relationship between ADHD diagnosis and behavioural variability and sleepiness ratings. The increased slow wave activity may represent a key neural mechanism underlying attentional fluctuations and challenges in ADHD, serving as a critical link between neurophysiological differences and behavioural performance across both clinical ADHD and neurotypical populations.

## Notes

### Competing Interest Statement

The authors have declared no competing interest.

### Funding

- National Health and Medical Research Council: APP2002454
- European Research Council: 101116748

