## Supplementary Figures and Table for "Sleep-like Slow Waves During Wakefulness Mediate Attention and Vigilance Difficulties in Adult Attention-Deficit/Hyperactivity Disorder"

**Supplementary Table**

**Supplementary Table A.**

*Demographics and Questionnaire Data*

| | ADHD (*N* = 32) | Neurotypical (*N* = 31) |
|---|---|---|
| **Adult ADHD Self-Report Scale (ASRS-5)** (mean (SD)) | 16.3 (2.9) | 7.8 (4.1) |
| **Wender Utah Rating Scale** (mean (SD)) | 52.4 (16.9) | |
| **Inventory of Depression and Anxiety Symptoms (IDAS-II)** (mean (SD)) | | |
| General Depression | 54.7 (12.6) | 39.1 (11.5) |
| Dysphoria | 28.1(7.8) | 18.0 (6.7) |
| Well Being | 20.8 (6.8) | 24.3 (8.2) |
| Panic | 13.9 (4.9) | 10.8 (3.3) |
| **Imaginal Process Inventory (IPI)** (mean (SD)) | 81.6 (14.2) | 63.8 (21.9) |
| **Morningness-Eveningness Questionnaire (MEQ)** (mean (SD)) | 40.8 (8.8) | 46.9 (7.6) |
| **Alcohol Use Disorders Identification Test** | 4.3 (3.4) | 3.0 (3.3) |
| **Drug Use Disorders Identification Test** | 1.9 (3.1) | 1.0 (2.4) |

***Note.*** ADHD = Attention Deficit/Hyperactivity Disorder. Higher scores on the ASRS-5 (total score) indicate greater ADHD symptom severity. Higher WURS scores reflect greater retrospectively reported childhood ADHD symptoms. Higher scores on the IDAS-II subscales reflect greater symptom severity within each domain. Higher scores on the IPI subset indicate more frequent daydreaming, vivid dream recall, and stronger immersion in internal experiences. Higher MEQ scores indicate a stronger morning preference, while lower scores reflect an evening preference. Higher AUDIT and DUDIT scores reflect greater alcohol consumption and drug use, respectively.

**Supplementary Table B.**

*ADHD Medication Use by DIVA Interview Diagnosis*

| Medications | Inattentive (*N* = 8) | Combined (*N* = 18) |
|---|---|---|
| **CNS Stimulants** | | |
| Methylphenidate | 3 (33.3%) | 7 (31.8%) |
| Dexamphetamine | 3 (33.3%) | 5 (22.7%) |
| Lisdexamfetamine | 1 (11.1%) | 7 (31.8%) |
| **Non-Stimulants** | | |
| Atomoxetine | 1 (11.1%) | 1 (4.5%) |

***Note***. ADHD = Attention Deficit/Hyperactivity Disorder. Medication categories are not mutually exclusive; the same individual may be counted for more than one medication. Percentages are the distribution across presentations. Note that one individual did not complete the DIVA interview but reported not taking any ADHD medication.

### Supplementary Figures

**Supp Figure 1.**

*Sustained Attention Response Task (SART) performance results 20s prior to a probe.*

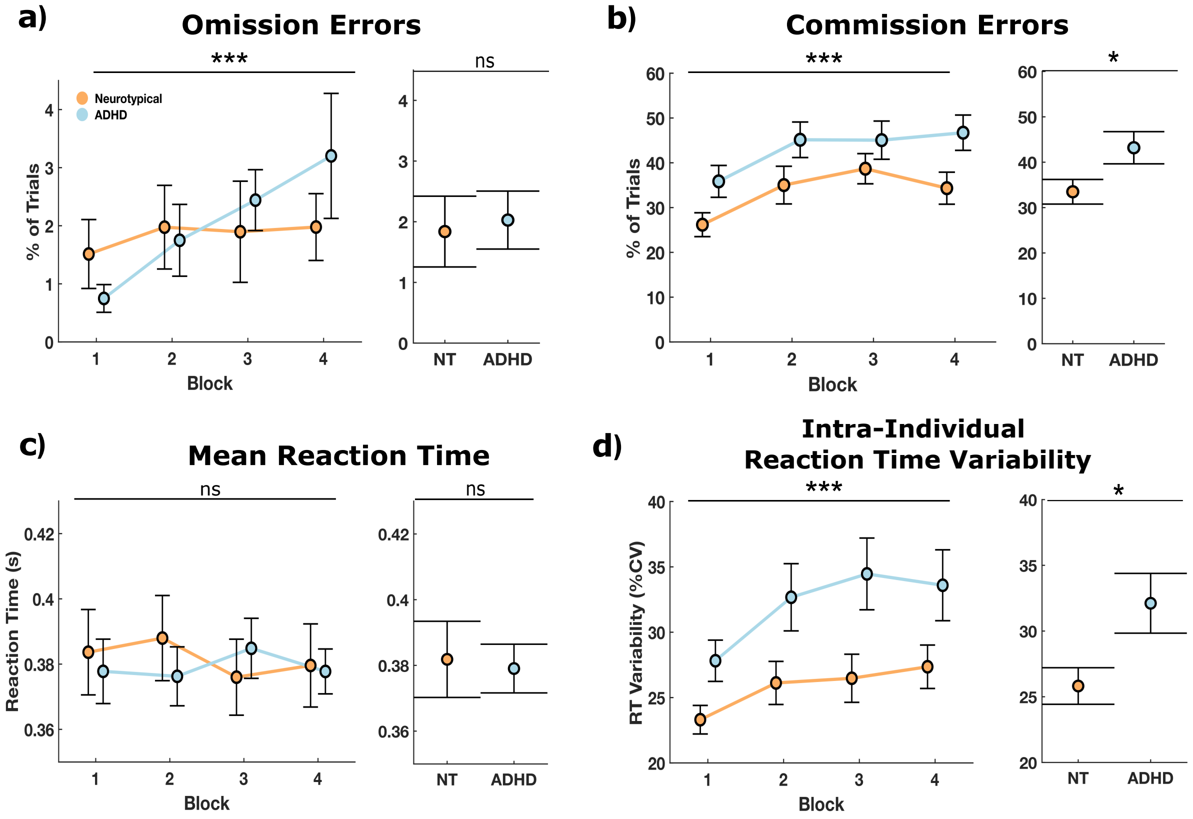


Performance metrics including: **a.** omission errors, **b.** commission errors, **c.** mean reaction time (RT), and **d.** RT variability (intra-individual coefficient of variation of reaction time). The left panels show the mean performance of each group (neurotypical (NT) in orange and ADHD in blue) across blocks (N = 4 blocks) with error bars representing the standard error of the performance variables. The right panels show the overall mean and standard error by group. Stars denote the significance level of block effect on performance (left panels) and performance differences between groups (right panels) as determined by linear mixed-effect models (ns: nonsignificant, *$p < 0.05$, **$p < 0.01$, ***$p < 0.001$). These results are consistent with whole task performance analysis.

**Supp Figure 2.**

*Sustained Attention Response Task (SART) performance and probe results by ADHD presentation (Combined n = 22, Inattentive n = 9).*

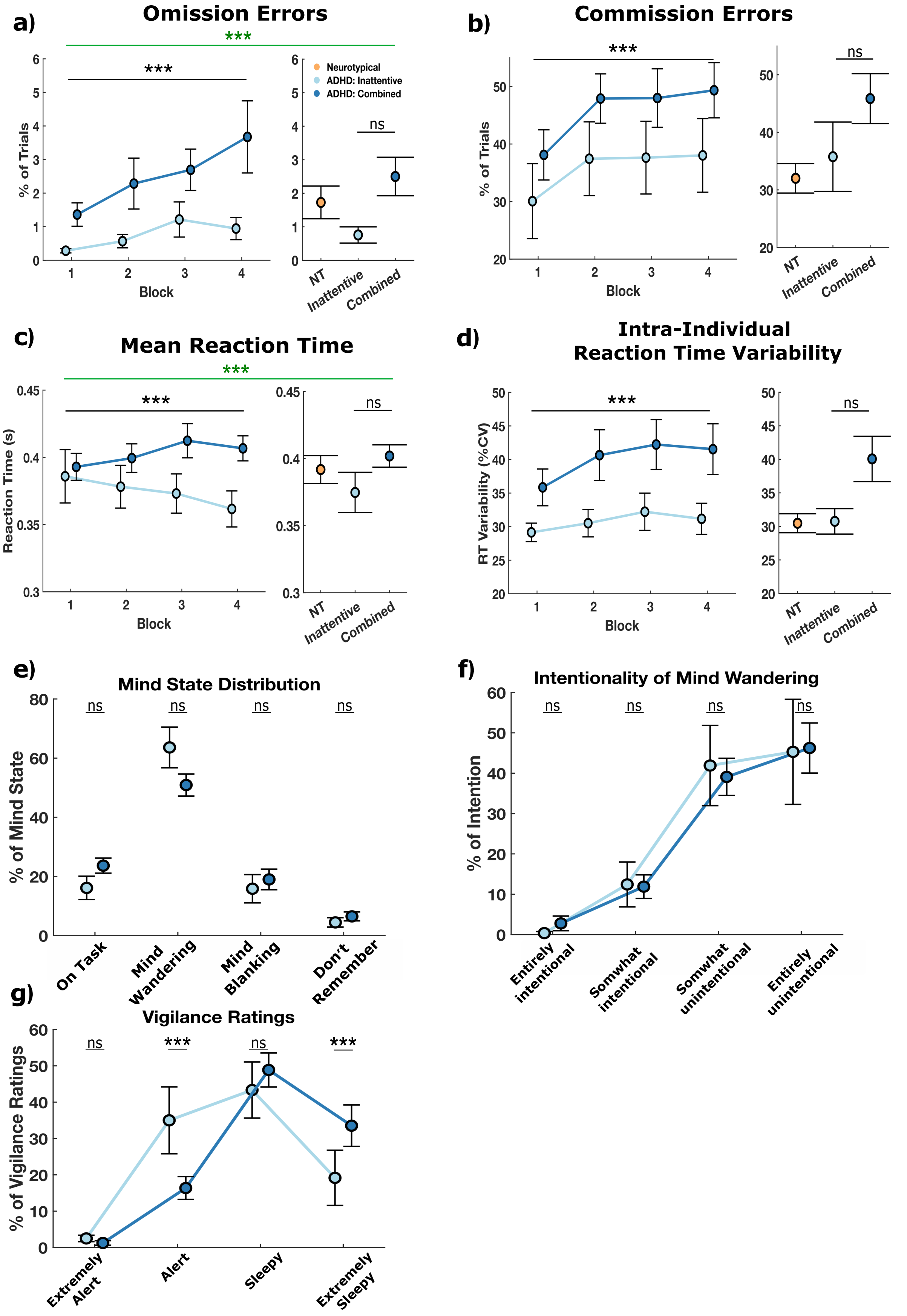


Performance metrics included: **a.** omissions, **b.** commissions, **c.** mean reaction time (RT), and **d.** RT variability (intra-individual coefficient of variation of reaction time). The left panels show the mean performance of each ADHD presentation (Inattentive in light blue and Combined in dark blue) across blocks (N = 4 blocks) with error bars representing the standard error (SE). The right panels show the overall mean and SE by presentation. Stars denote the significance level of block on performance (left panels), performance differences between presentation (right panels) and the presentation-by-block interaction (green stars) as determined by linear mixed-effect models (ns: nonsignificant, *$p < 0.05$, **$p < 0.01$, ***$p < 0.001$). Panels **e.**, **f.**, and **g.** show probe results. Stars denote significant levels of presentation differences per category.
